# The potential role of viruses controlling phytoplankton community size structure

**DOI:** 10.64898/2026.03.03.709231

**Authors:** Nazia Mojib, Xabier Irigoien

## Abstract

The size structure of phytoplankton communities plays a key role in the fate of carbon fixed by photosynthesis. Whether phytoplankton cells sink, enter the microbial loop, or are consumed by larger organisms is generally determined by their size. Grazing has been advanced as a factor determining size structure, but sources of mortality other than grazing, such as viruses also are recognized to be important. Based on the observation that cell size and genome size are related in phytoplankton, we hypothesize that viruses can also play a role in shaping the size structure of the phytoplankton community. Because cell size is related to genome size, we suggest that phytoplankton species with larger genomes will have a more developed immune system to defend against viral infection. As a first step to test this hypothesis, we screened the published transcriptomes of 125 phytoplankton species for expressed viral and immune-response related genes. We found a significant negative correlation between host-cell size and viral-gene diversity, and a positive correlation between host-cell size and the number of immune-response related genes. Our hypothesis supported by preliminary findings opens new pathways to explore whether we should consider viruses as an additional evolutionary driver for larger phytoplankton size, along with grazing and nutrients.

## Introduction

Phytoplankton account for about 40 % of the photosynthesis (Falkowski, 1994) on earth and consequently play a key role in biogeochemical cycles (Falkowski *et al*., 2000). Phytoplankton blooms, in particular, function by regulating the carbon fluxes in marine ecosystems (Behrenfeld & Boss, 2014). During blooms, phytoplankton biomass rapidly increases, by orders of magnitude, affecting different traits of ecosystem function - from phenology to biogeochemical cycles (Behrenfeld & Boss, 2014). Phytoplankton blooms are dominated by large cells (Irigoien *et al*., 2004) and are suggested to be a biomass accumulation of species with lower mortality (Behrenfeld & Boss, 2014) because of their size and/or defensive structures, such as spines, which may reduce grazing pressure (Irigoien *et al*., 2005). Viruses are also a major source of phytoplankton mortality (Brussaard, 2004, Fuhrman, 1999, Suttle, 2005, Suttle *et al*., 1990, Weitz & Wilhelm, 2012). Although cell size itself is not directly linked to the success of viral infection, it is strongly related to genome size (Holm-Hansen, 1969); in fact, evidence shows a relationship between genome size and gene number in unicellular eukaryotic organisms (Hou & Lin, 2009, Lynch, 2006). Therefore, we hypothesize that larger cells are likely to contain more genetic resources with the potential to develop more efficient immune responses against viral infection and that viruses would also contribute to the size structure of the phytoplankton blooms.

## Material and Methods

### Selection of phytoplankton species

From the 671 transcriptomes with 306 unique species (as of 18 December 2014) available from the Marine Microbial Eukaryote Transcriptome Sequencing Project (MMETSP:https://zenodo.org/records/3247846)) (Keeling *et al*., 2014), we selected 125 free-living autotroph planktonic organisms with unique species names and known cell sizes/volumes for RNA-Seq analysis (Table I).

### Detection of viruses in the transcriptome of phytoplankton

The raw Illumina RNA-Seq data in fastq format of 125 phytoplankton species were obtained from CAMERA’s (Community cyberinfrastructure for Advanced Microbial Ecology Research and Analysis) MMETSP portal (http://camera.calit2.net/mmetsp/) (Sun *et al*., 2011) (current data as of 18 December 2014) under the unique identifiers, as shown in Table I. All the datasets can now be retrieved from Zenodo: https://zenodo.org/records/3247846. The raw reads were prefiltered for Illumina primers/adaptors, control DNA (phiX174), and 50-bp long with Phred score ≥ 33. The RNA-Seq analysis for the viral-gene expression of each phytoplankton species was performed using CLC Genomics Workbench v.7.5. now available on QIAGEN Digital Insights. The reads were aligned to the viral genomes/genes (n= 5,401/188,972) downloaded on 29 May 2014 from the viral genomes resource (National Center for Biotechnology Information) (Supplementary Data 1). The RNA-Seq analysis was performed for each species using the following parameters: the numbers of mismatches allowed were two, a minimum length fraction of 0.8, a minimum similarity fraction of 0.8, and a maximum hit for a read of 1. Only the genes aligned with > 50 reads were considered as expressed (Supplementary Table SII). Cell volumes of each phytoplankton species were deduced from the sources as indicated (Table I).

### Detection of immune-response-related genes

The first step was to prepare a reference database of immune-response-related genes (IRGs). For this, all plant IRGs were extracted from *Arabidopsis* and *Manihotesculenta* (cassava) as identified by Leal et al.(Leal *et al*., 2013) Then, the homologous sequences of IRGs related to these plants were searched from the genomes of the model diatoms, *Phaeodactylum tricornutum* CCAP 1055/1 (NZ_ABQD00000000) (Bowler *et al*., 2008) and *Thalassiosira pseudonana* CCMP1335 (NZ_AAFD00000000) (Armbrust *et al*., 2004), using Blastn. All the homologous IRGs from both diatoms (n=991) were then combined for use as a reference (Supplementary Data 2) for screening the 125-tested phytoplankton species; their abundance was inferred similar to the methods described for viral gene detection (Supplementary Table SII).

## Results and discussion: Correlation among host-cell size, its immune related and expressed viral genes

We use a direct approach to detect and assess the diversity of intra-host-expressed viral genes by aligning the RNA-Seq reads of each phytoplankton species to a collection of viral genomes/genes (n= 5,401/188,972). We searched for homologous IRGs in each phytoplankton species using the in-house-generated reference database (n=991) for IRGs created from the genomes of two model diatoms, *Phaeodactylum tricornutum* and *Thalassiosira pseudonana* (Supplementary Data 1, 2&3). Viruses present in each phytoplankton species were assessed by counting the number of genes aligned to 50 or more unique reads (Supplementary Table SI, Supplementary Data 3). Similarly, the IRGs expressed in each species were assessed according to the number of homologous genes that aligned to reference IRGs (Supplementary Table SII). Using previously estimated cell volumes for each phytoplankton species (Table I), we found a significantly negative relationship between cell volume and expressed viral transcript diversity (viral diversity = -15.32 ln (cell volume) + 312.49, r^2^=0.15, n=125, p<0.001, Fig. 1a), and a significantly positive relationship between cell volume and the number of IRGs (immune related genes = 14.83 ln (cell volume) + 159.02, r^2^=0.16, p<0.001, n=125, Fig. 1b). There was no significantly negative relationship between host-cell size and number of detected viruses (r^2^ = 0.03) when viral numbers included minimally expressed genes (i.e., mapped unique reads between 0 and 10), but when moderately to highly expressed genes (i.e., unique reads greater than 50) were included in the analysis a significantly negative relationship (r^2^=0.15, 0.14) was observed (Fig. 2). This result suggests that the number of viral species is independent of host-cell size, but that the quantity of viral genes expressed is limited by host-cell size. Presumably, larger host cells, most likely with a larger genome, have greater immune-gene content and expression, which limits the expression of viral genes. Although significant, both relationships show high-level dispersion, where only 15 to 16 % of the variance is explained by size.

**Figure 1.**
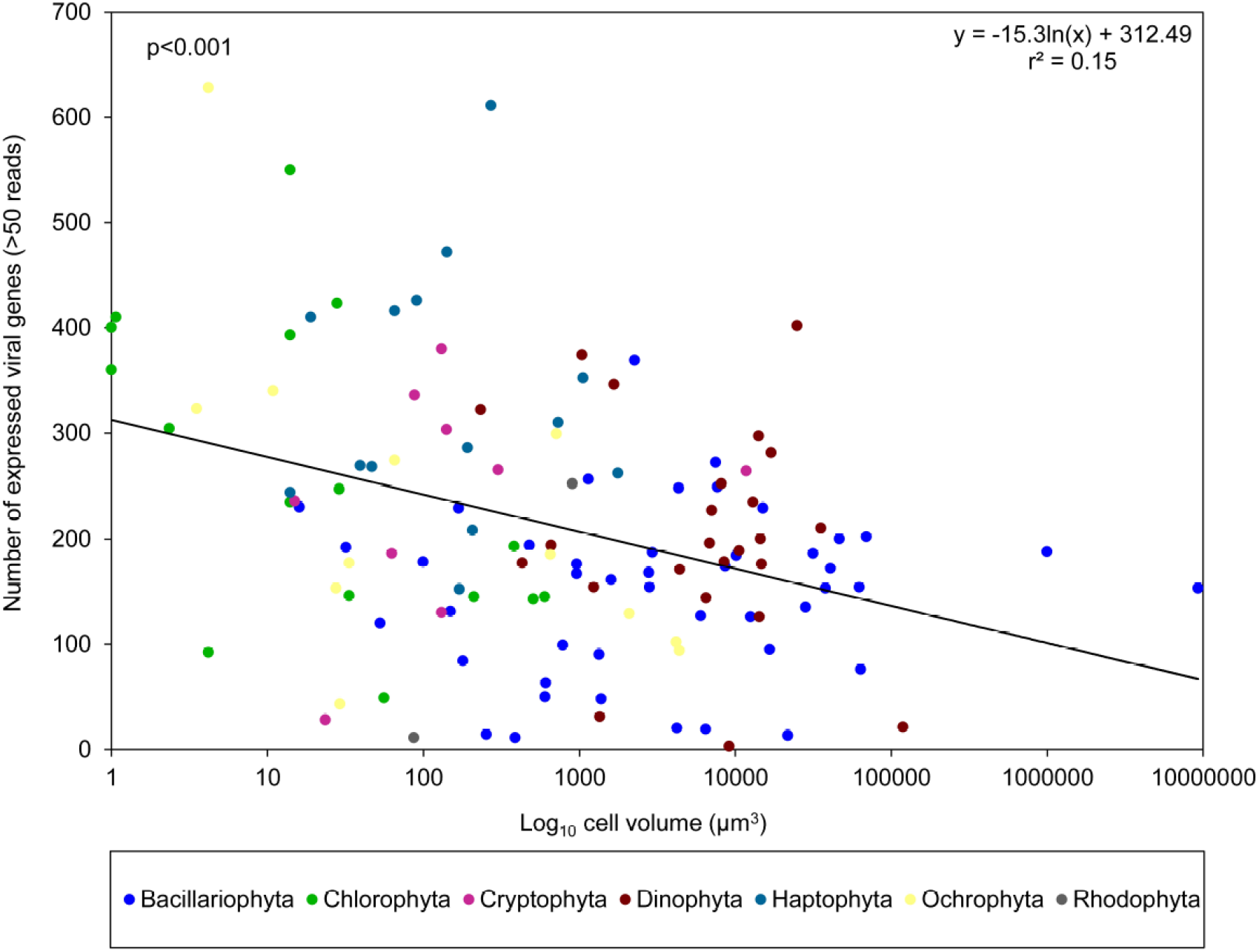
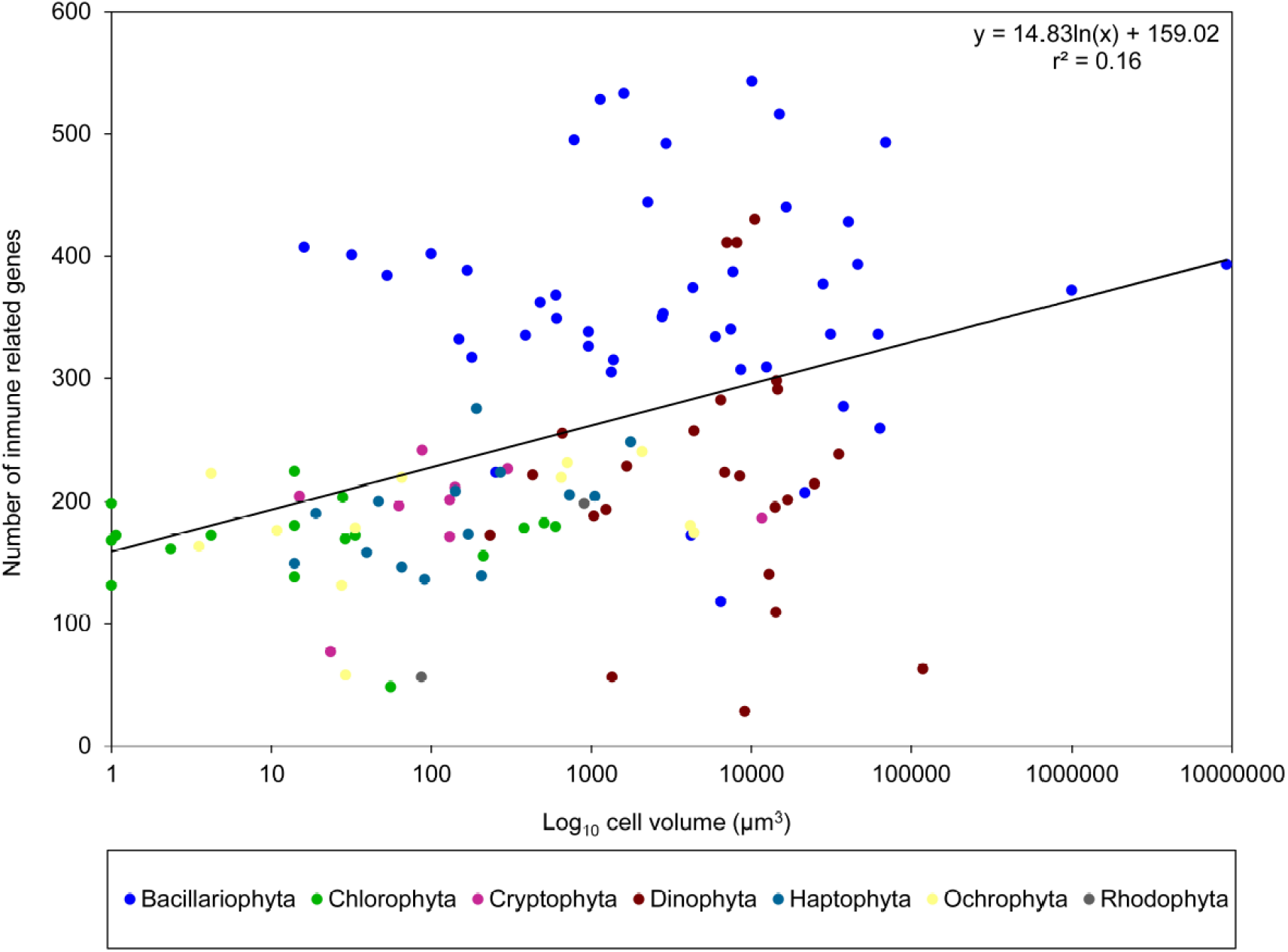
The relationship of viruses and immune-related genes (IRGs) with phytoplankton size. (a) A scatterplot showing negative correlation between RNA-Seq-derived number of viruses detected (at > 50 unique-mapped reads) in phytoplankton species (n=125) (Supplementary Table SI) and their cell volumes (Table I). (b) A scatterplot showing positive correlation between RNA-Seq-derived number of homologous immune-related genes in 125 phytoplankton species (Supplementary Table SII) and their cell volumes.

**Figure 2.**
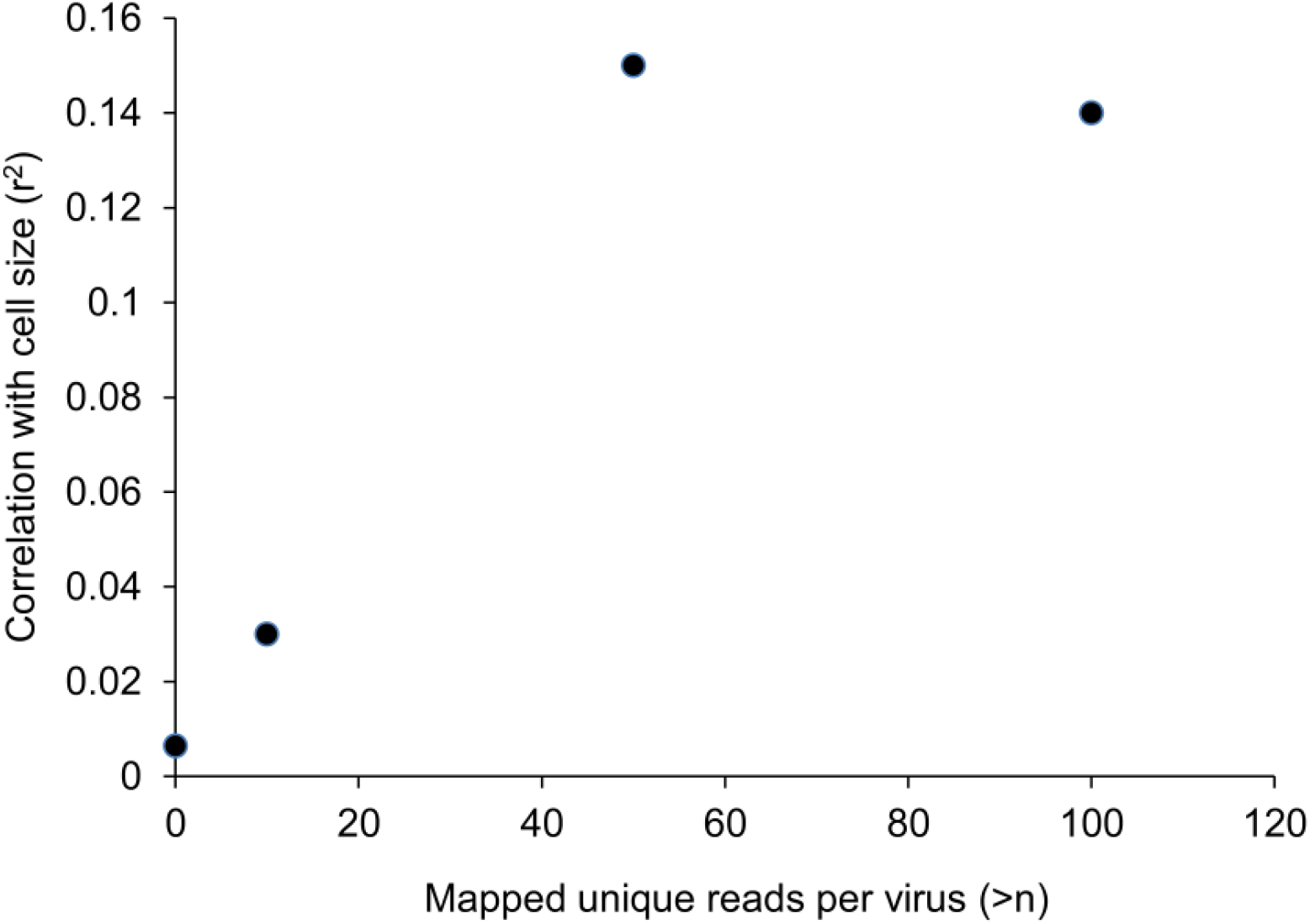
The scatter plot of correlation coefficient (r^2^) depicting relationship between host-cell size and number of detected viruses against mapped unique reads per virus. There was no significantly negative relationship between host-cell size and number of detected viruses (r^2^ = 0.03) when mapped unique reads were between 0 and 10, but when unique reads greater than 50 were included in the analysis a significant negative relationship (r^2^=0.15, 0.14) was observed.

Because our knowledge on the viruses that infect phytoplankton and the immune mechanisms of phytoplankton is still limited (Gregory *et al*., 2019), our measurements of expressed viral diversity and quantity of IRGs can only be considered rough estimates. Furthermore, some groups included in our analysis, such as dinoflagellates, contain huge, variable genomes that are difficult to understand (Hackett *et al*., 2004). Due to a lack of information on immune mechanisms in phytoplankton, we measured IRG expression by comparison with genes in the existing genome sequences of two model diatoms, *P. tricornutum* and *T. pseudonana*, which are homologous to IRGs identified in plants (*Arabidopsis thaliana*). Therefore, the calculated number of expressed IRGs is biased towards groups that are evolutionarily close to these diatoms; for this reason, tested diatoms (Bacillariophyta) (Fig. 1b) showed the maximum homology with the reference IRGs. The relationship between cell size and DNA content is general (Cavalier-Smith, 1985), however, to what extent a larger genome entails higher complexity (the C-paradox) has long been a subject of debate (Gregory, 2001). The advent of genomic sequencing has made studying the relationship between genome size and number of genes more attainable; for example, in unicellular eukaryotes there is a significant relationship between haploid genome size and number of genes (Lynch, 2006). Therefore, it is reasonable to infer that larger phytoplankton cells may have more genes allocated to the immune system that would result in lower rates of infection, as suggested by our results. A larger genome also results in faster genomic evolution (Connolly *et al*., 2008, Oliver *et al*., 2007), an advantage in the coevolution of the arms race between viruses and eukaryotic cells (Smetacek, 2001). The advantages for a larger-sized genome may, therefore, be two-fold: the immediate advantage of a more sophisticated immune system during favorable growing conditions, which will allow the species to bloom and an evolutionary advantage in the context of the Red Queen’s race with viruses.

Finally, viruses playing a role in the size structure of phytoplankton blooms do not exclude other factors such as grazing and nutrients (Irigoien *et al*., 2005). By investigating the types of viruses among phytoplankton species, similar to previous work (Monier *et al*., 2008), we found that a significant portion of the viruses expressed belong to the giant virus family. In fact, dsDNA large viruses dominate the top 10 viruses detected in all species evaluated (Fig.3), suggesting their predominance in marine phytoplankton species in general.

**Figure 3.**
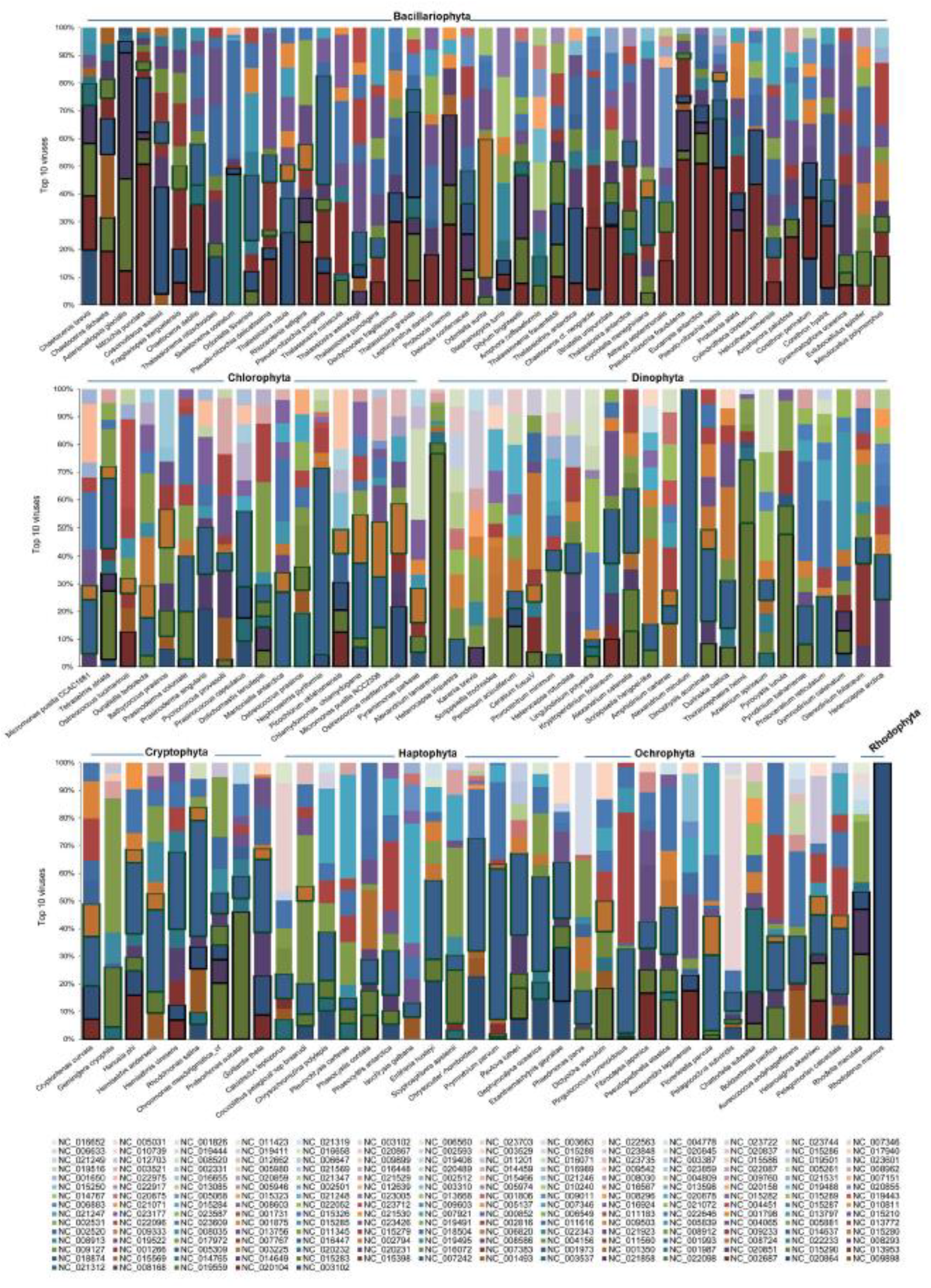
Top 10 viruses detected in all species evaluated

## Concluding remarks

The roles of grazing and nutrients as evolutionary driving forces towards larger sizes in phytoplankton are a subject of debate (Finkel *et al*., 2007, Irigoien *et al*., 2005, Litchman *et al*., 2009); however, our results indicate that viruses may also play a role and should, therefore, be included in the debate. In fact, viruses impact the evolution of eukaryotic cells at different levels (Baluska, 2009, Rohwer *et al*., 2009, Rohwer & Thurber, 2009), which needs to be considered in the context of phytoplankton-size evolution. In general, we expect that larger unicellular eukaryotes, which have larger genomes that often have a more number of genes, should be considered in the evolutionary size argument. A larger number of genes provides these larger cells with traits and capacities beyond those directly related to size such as surface/volume effects. Here we highlight the relationship between phytoplankton cell size and virus resistance, but other aspects such as production of allelopathic compounds and toxins also need to be considered in the evolutionary size debate.

## Supporting information

Supplementary Table SI. Screening results of 125 RNA-Seq libraries for intra host expressed viral genes.

Supplementary Table SII. Screening results of 125 RNA-Seq libraries for immune related genes (IRGs).

List of 125 phytoplankton species with their MMETSP identifiers and cell volumes along with the references.

## Acknowledgements

We thank the Gordon and Betty Moore Foundation, the National Center for Genome Resources, the National Center for Marine Algae and Microbiota, and all the participants of The Marine Microbial Eukaryote Transcriptome Sequencing Project for making the data publicly available. This research was supported by baseline funding provided by King Abdullah University of Science and Technology to Prof. Xabier Irigoien.

## Author contributions

N.M. and X.I. designed the study; N.M. performed the analysis; X.I. and N.M. wrote the paper.

