## Supplementary Table SI. Screening results of 125 RNA-Seq libraries for intra host expressed viral genes. for "The potential role of viruses controlling phytoplankton community size structure"

| MMETSP ID | Phylum | Species | Total library size (reads) | Mapped reads to virus database | %Total viral reads mapped |
|---|---|---|---|---|---|
| MMETSP0316 | Bacillariophyta | *Amphora coffeaeformis* | 63726418 | 48882 | 0.08 |
| MMETSP0705 | Bacillariophyta | *Asterionellopsis glacialis* | 36609940 | 37372 | 0.1 |
| MMETSP1449 | Bacillariophyta | *Attheya septentrionalis* | 68907220 | 68465 | 0.1 |
| MMETSP1435 | Bacillariophyta | *Chaetoceros brevis* | 18446420 | 12997 | 0.07 |
| MMETSP1336 | Bacillariophyta | *Chaetoceros cf. neogracile* | 29886328 | 47647 | 0.16 |
| MMETSP0150 | Bacillariophyta | *Chaetoceros debilis* | 51515312 | 77782 | 0.15 |
| MMETSP1447 | Bacillariophyta | *Chaetoceros dichaeta* | 44854282 | 24768 | 0.06 |
| MMETSP1066 | Bacillariophyta | *Coscinodiscus wailesii* | 66050422 | 63259 | 0.1 |
| MMETSP1057 | Bacillariophyta | *Cyclotella meneghiniana* | 84637736 | 58794 | 0.07 |
| MMETSP0017_2 | Bacillariophyta | *Cylindrotheca closterium* | 34126456 | 12259 | 0.04 |
| MMETSP0580 | Bacillariophyta | *Dactyliosolen fragilissimus* | 39877006 | 60960 | 0.15 |
| MMETSP1058 | Bacillariophyta | *Detonula confervacea* | 66235192 | 61607 | 0.09 |
| MMETSP0998 | Bacillariophyta | *Ditylum brightwellii* | 62798858 | 79646 | 0.13 |
| MMETSP1437 | Bacillariophyta | *Eucampia antarctica* | 21265386 | 29151 | 0.14 |
| MMETSP0733 | Bacillariophyta | *Fragilariopsis kerguelensis* | 49019016 | 164496 | 0.34 |
| MMETSP1171 | Bacillariophyta | *Helicotheca tamensis* | 39953914 | 145813 | 0.36 |
| MMETSP1362 | Bacillariophyta | *Leptocylindrus danicus* | 39861406 | 47065 | 0.12 |
| MMETSP0744 | Bacillariophyta | *Nitzschia punctata* | 90547632 | 123724 | 0.14 |
| MMETSP0015_2 | Bacillariophyta | *Odontella aurita* | 42084656 | 8985 | 0.02 |
| MMETSP0160_2 | Bacillariophyta | *Odontella Sinensis* | 49162656 | 73252 | 0.15 |
| MMETSP0174 | Bacillariophyta | *Proboscia alata* | 53939606 | 54587 | 0.1 |
| MMETSP0816 | Bacillariophyta | *Proboscia inermis* | 51566466 | 83454 | 0.16 |
| MMETSP0327 | Bacillariophyta | *Pseudo-nitzschia delicatissima* | 76463666 | 87170 | 0.11 |
| MMETSP0850 | Bacillariophyta | *Pseudo-nitzschia fraudulenta* | 35381966 | 52843 | 0.15 |
| MMETSP1423 | Bacillariophyta | *Pseudo-nitzschia heimii* | 34362058 | 63038 | 0.18 |
| MMETSP1060 | Bacillariophyta | *Pseudo-nitzschia pungens* | 75981556 | 196613 | 0.26 |
| MMETSP0789 | Bacillariophyta | *Rhizosolenia setigera* | 53486746 | 98175 | 0.18 |
| MMETSP0013_2 | Bacillariophyta | *Skeletonema costatum* | 36792038 | 6116 | 0.02 |
| MMETSP0794_2 | Bacillariophyta | *Stephanopyxis turris* | 28656286 | 64707 | 0.23 |
| MMETSP0800 | Bacillariophyta | *Striatella unipunctata* | 48336382 | 90171 | 0.19 |
| MMETSP0786 | Bacillariophyta | *Thalassionema frauenfeldii* | 51370380 | 70668 | 0.14 |
| MMETSP0156 | Bacillariophyta | *Thalassionema nitzschioides* | 52529482 | 77421 | 0.15 |
| MMETSP0902 | Bacillariophyta | *Thalassiosira antarctica* | 32870602 | 39903 | 0.12 |
| MMETSP0492 | Bacillariophyta | *Thalassiosira gravida* | 39941506 | 33097 | 0.08 |
| MMETSP0737 | Bacillariophyta | *Thalassiosira miniscula* | 72933184 | 116041 | 0.16 |
| MMETSP1067 | Bacillariophyta | *Thalassiosira punctigera* | 56431354 | 63952 | 0.11 |
| MMETSP0403 | Bacillariophyta | *Thalassiosira rotula* | 80967272 | 87572 | 0.11 |
| MMETSP0878 | Bacillariophyta | *Thalassiosira weissflogii* | 52047092 | 84549 | 0.16 |
| MMETSP0152 | Bacillariophyta | *Thalassiothrix antarctica* | 45679326 | 103521 | 0.23 |
| MMETSP1065 | Bacillariophyta | *Amphiprora paludosa* | 66002034 | 52236 | 0.08 |
| MMETSP0169 | Bacillariophyta | *Corethron pennatum* | 51460954 | 74697 | 0.15 |
| MMETSP0010_2 | Bacillariophyta | *Corethron hystrix* | 23183502 | 5707 | 0.02 |
| MMETSP0009_2 | Bacillariophyta | *Grammatophora oceanica* | 31641830 | 41344 | 0.13 |

This file has been converted from its original format for security purposes. Please use C6747616631EE as a reference.

| | | | | | |
|---|---|---|---|---|---|
| MMETSP0696 | Bacillariophyta | *Extubocellulus spinifer* | 44300740 | 57296 | 0.13 |
| MMETSP1070 | Bacillariophyta | *Minutocellus polymorphus* | 69826230 | 91474 | 0.13 |
| MMETSP1460 | Chlorophyta | *Bathycoccus prasinos* | 55777388 | 154120 | 0.28 |
| MMETSP1392 | Chlorophyta | *Chlamydomonas chlamydogama* | 64172020 | 77416 | 0.12 |
| MMETSP0033_2 | Chlorophyta | *Dolichomastix tenuilepis* | 55347526 | 206376 | 0.37 |
| MMETSP1126 | Chlorophyta | *Dunaliella tertiolecta* | 58248576 | 97324 | 0.17 |
| MMETSP1106 | Chlorophyta | *Mantoniella antarctica* | 48679686 | 92871 | 0.19 |
| MMETSP1401 | Chlorophyta | *Micromonas pusilla CCAC1681* | 43995616 | 147094 | 0.33 |
| MMETSP1327 | Chlorophyta | *Micromonas pusilla RCC2306* | 45830388 | 99415 | 0.22 |
| MMETSP0034_2 | Chlorophyta | *Nephroselmis pyriformis* | 46610224 | 56934 | 0.12 |
| MMETSP0939 | Chlorophyta | *Ostreococcus lucimarinus* | 62631002 | 154734 | 0.25 |
| MMETSP0929 | Chlorophyta | *Ostreococcus mediterraneus* | 87232616 | 169210 | 0.19 |
| MMETSP0933 | Chlorophyta | *Ostreococcus prasinos* | 54576496 | 106378 | 0.19 |
| MMETSP1161 | Chlorophyta | *Picochlorum oklahomensis* | 41993614 | 33646 | 0.08 |
| MMETSP0941 | Chlorophyta | *Prasinococcus capsulatus* | 47027436 | 53415 | 0.11 |
| MMETSP0806_2 | Chlorophyta | *Prasinoderma coloniale* | 52641318 | 173789 | 0.33 |
| MMETSP1315 | Chlorophyta | *Prasinoderma singularis* | 45156550 | 143568 | 0.32 |
| MMETSP1316 | Chlorophyta | *Pycnococcus provasolii* | 41226934 | 115557 | 0.28 |
| MMETSP0819 | Chlorophyta | *Tetraselmis striata* | 40593864 | 96396 | 0.24 |
| MMETSP0058 | Chlorophyta | *Pyramimonas parkeae* | 67915958 | 66592 | 0.1 |
| MMETSP1050 | Cryptophyta | *Cryptomonas curvata* | 56053624 | 203478 | 0.36 |
| MMETSP0799 | Cryptophyta | *Geminigera cryophila* | 58337084 | 822739 | 1.41 |
| MMETSP1048 | Cryptophyta | *Hanusia phi* | 52458676 | 116212 | 0.22 |
| MMETSP0043_2 | Cryptophyta | *Hemiselmis andersenii* | 78445892 | 139123 | 0.18 |
| MMETSP1356 | Cryptophyta | *Hemiselmis viresens* | 52614122 | 110593 | 0.21 |
| MMETSP1047 | Cryptophyta | *Rhodomonas salina* | 54321504 | 267760 | 0.49 |
| MMETSP0047_2 | Cryptophyta | *Chroomonas mesostigmatica_cf* | 67406484 | 315864 | 0.47 |
| MMETSP1049 | Cryptophyta | *Proteomonas sulcata* | 48515622 | 85679 | 0.18 |
| MMETSP0046_2 | Cryptophyta | *Guillardia theta* | 42158888 | 57787 | 0.14 |
| MMETSP0790 | Dinophyta | *Alexandrium catenella* | 42794264 | 85833 | 0.2 |
| MMETSP0328 | Dinophyta | *Alexandrium minutum* | 22624640 | 5614 | 0.02 |
| MMETSP0378 | Dinophyta | *Alexandrium tamarense* | 28696260 | 123307 | 0.43 |
| MMETSP0258 | Dinophyta | *Amphidinium carterae* | 52247898 | 129156 | 0.25 |
| MMETSP1074 | Dinophyta | *Ceratium fususV* | 54627688 | 73605 | 0.13 |
| MMETSP0797 | Dinophyta | *Dinophysis acuminata* | 48685084 | 104571 | 0.21 |
| MMETSP0116_2 | Dinophyta | *Durinskia baltica* | 46992432 | 88163 | 0.19 |
| MMETSP0503 | Dinophyta | *Heterocapsa rotundata* | 43256526 | 127293 | 0.29 |
| MMETSP0448 | Dinophyta | *Heterocapsa triquestra* | 45359332 | 126875 | 0.28 |
| MMETSP0027 | Dinophyta | *Karenia brevis* | 26748951 | 48735 | 0.18 |
| MMETSP0120_2 | Dinophyta | *Kryptoperidinium foliaceum* | 63629800 | 89468 | 0.14 |
| MMETSP1032 | Dinophyta | *Lingulodinium polyedra* | 65984202 | 218632 | 0.33 |
| MMETSP0370 | Dinophyta | *Peridinium aciculiferum* | 41244260 | 98586 | 0.24 |
| MMETSP0267 | Dinophyta | *Prorocentrum minimum* | 25887799 | 68794 | 0.27 |
| MMETSP0367 | Dinophyta | *Scrippsiella hangoei-like* | 46219246 | 92290 | 0.2 |
| MMETSP0270 | Dinophyta | *Scrippsiella trochoidea* | 36147274 | 82914 | 0.23 |
| MMETSP0229_2 | Dinophyta | *Pyrocystis lunula* | 62650660 | 18698 | 0.03 |
| MMETSP0796 | Dinophyta | *Pyrodinium bahamense* | 62709420 | 104562 | 0.17 |

This file has been converted from its original format for security purposes. Please use C6747616631EE as a reference.

| | | | | | |
|---|---|---|---|---|---|
| MMETSP0228 | Dinophyta | *Protoceratium reticulatum* | 46869592 | 155637 | 0.33 |
| MMETSP0784 | Dinophyta | *Gymnodinium catenatum* | 73471400 | 88772 | 0.12 |
| MMETSP0118_2 | Dinophyta | *Glenodinium foliaceum* | 45761730 | 97671 | 0.21 |
| MMETSP0225 | Dinophyta | *Thoracosphaera heimii* | 54872800 | 44742 | 0.08 |
| MMETSP1036_2 | Dinophyta | *Azadinium spinosum* | 51318154 | 82412 | 0.16 |
| MMETSP1441 | Dinophyta | *Heterocapsa arctica* | 54674410 | 188914 | 0.35 |
| MMETSP1334 | Haptophyta | *Calcidiscus leptoporus* | 44404398 | 223589 | 0.5 |
| MMETSP0143 | Haptophyta | *Chrysochromulina polylepis* | 74214716 | 230606 | 0.31 |
| MMETSP0164_2 | Haptophyta | *Coccolithus pelagicus ssp braarudi* | 53235070 | 220296 | 0.41 |
| MMETSP0994 | Haptophyta | *Emiliania huxleyi* | 25066298 | 102303 | 0.41 |
| MMETSP0944 | Haptophyta | *Isochrysis galbana* | 45534184 | 117196 | 0.26 |
| MMETSP1444 | Haptophyta | *Phaeocystis antarctica* | 47646454 | 238498 | 0.5 |
| MMETSP1465 | Haptophyta | *Phaeocystis cordata* | 38854414 | 107708 | 0.28 |
| MMETSP1136 | Haptophyta | *Pleurochrysis carterae* | 43469752 | 182235 | 0.42 |
| MMETSP1333 | Haptophyta | *Scyphosphaera apsteinii* | 40529618 | 179856 | 0.44 |
| MMETSP1335 | Haptophyta | *Chrysoculter rhomboideus* | 43272560 | 173265 | 0.4 |
| MMETSP0006_2 | Haptophyta | *Prymnesium parvum* | 42932420 | 101168 | 0.24 |
| MMETSP1363 | Haptophyta | *Gephyrocapsa oceanica* | 54668892 | 193881 | 0.35 |
| MMETSP1463 | Haptophyta | *Pavlova lutheri* | 45354612 | 161158 | 0.36 |
| MMETSP1464 | Haptophyta | *Exanthemachrysis gayraliae* | 48832876 | 178282 | 0.37 |
| MMETSP0915 | Ochrophyta | *Aureococcus anophagefferens* | 46827590 | 233634 | 0.5 |
| MMETSP1319 | Ochrophyta | *Bolidomonas pacifica* | 43287000 | 168398 | 0.39 |
| MMETSP0292 | Ochrophyta | *Heterosigma akashiwo* | 32622070 | 67692 | 0.21 |
| MMETSP0888 | Ochrophyta | *Pelagomonas calceolata* | 42273462 | 114910 | 0.27 |
| MMETSP1339 | Ochrophyta | *Fibrocapsa japonica* | 51008030 | 95992 | 0.19 |
| MMETSP1174 | Ochrophyta | *Dictyocha speculum* | 41736862 | 55971 | 0.13 |
| MMETSP1068 | Ochrophyta | *Pseudopedinella elastica* | 59467734 | 100636 | 0.71 |
| MMETSP0947 | Ochrophyta | *Chattonella subsalsa* | 75999072 | 122431 | 0.16 |
| MMETSP1160 | Ochrophyta | *Pinguiococcus pyrenoidosus* | 60971594 | 239881 | 0.39 |
| MMETSP1323 | Ochrophyta | *Florenciella parvula* | 24016606 | 116414 | 0.48 |
| MMETSP1163 | Ochrophyta | *Phaeomonas parva* | 47821364 | 108177 | 0.23 |
| MMETSP0890 | Ochrophyta | *Aureoumbra lagunensis* | 24957712 | 144207 | 0.58 |
| MMETSP0882 | Ochrophyta | *Pelagococcus subviridis* | 36185352 | 153917 | 0.43 |
| MMETSP0167 | Rhodophyta | *Rhodella maculata* | 54377336 | 135000 | 0.25 |
| MMETSP0011_2 | Rhodophyta | *Rhodosorus marinus* | 42420550 | 5790 | 0.01 |

This file has been converted from its original format for security purposes. Please use C6747616631EE as a reference.
