## Supplementary Table SII. Screening results of 125 RNA-Seq libraries for immune related genes (IRGs). for "The potential role of viruses controlling phytoplankton community size structure"

| MMETSP ID | Phylum | Species | %Total reads mapped | Total library size (reads) | Mapped reads to IRGs |
|---|---|---|---|---|---|
| MMETSP0316 | Bacillariophyta | *Amphora coffeaeformis* | 0.06 | 63726418 | 41049 |
| MMETSP0705 | Bacillariophyta | *Asterionellopsis glacialis* | 0.04 | 36609940 | 13162 |
| MMETSP1449 | Bacillariophyta | *Attheya septentrionalis* | 0.17 | 68907220 | 119607 |
| MMETSP1435 | Bacillariophyta | *Chaetoceros brevis* | 0.11 | 18446420 | 19967 |
| MMETSP1336 | Bacillariophyta | *Chaetoceros cf. neogracile* | 0.22 | 29886328 | 65929 |
| MMETSP0150 | Bacillariophyta | *Chaetoceros debilis* | 0.24 | 51515312 | 123720 |
| MMETSP1447 | Bacillariophyta | *Chaetoceros dichaeta* | 0.21 | 44854282 | 94959 |
| MMETSP1066 | Bacillariophyta | *Coscinodiscus wailesii* | 0.1 | 66050422 | 65097 |
| MMETSP1057 | Bacillariophyta | *Cyclotella meneghiniana* | 0.27 | 84637736 | 229104 |
| MMETSP0017_2 | Bacillariophyta | *Cylindrotheca closterium* | 0.24 | 34126456 | 81384 |
| MMETSP0580 | Bacillariophyta | *Dactyliosolen fragilissimus* | 0.1 | 39877006 | 39971 |
| MMETSP1058 | Bacillariophyta | *Detonula confervacea* | 0.31 | 66235192 | 207515 |
| MMETSP0998 | Bacillariophyta | *Ditylum brightwellii* | 0.2 | 62798858 | 122630 |
| MMETSP1437 | Bacillariophyta | *Eucampia antarctica* | 0.11 | 21265386 | 22811 |
| MMETSP0733 | Bacillariophyta | *Fragilariopsis kerguelensis* | 0.1 | 49019016 | 47530 |
| MMETSP1171 | Bacillariophyta | *Helicotheca tamensis* | 0.11 | 39953914 | 42603 |
| MMETSP1362 | Bacillariophyta | *Leptocylindrus danicus* | 0.09 | 39861406 | 36952 |
| MMETSP0744 | Bacillariophyta | *Nitzschia punctata* | 0.07 | 90547632 | 60993 |
| MMETSP0015_2 | Bacillariophyta | *Odontella aurita* | 0.23 | 42084656 | 98102 |
| MMETSP0160_2 | Bacillariophyta | *Odontella Sinensis* | 0.15 | 49162656 | 72255 |
| MMETSP0174 | Bacillariophyta | *Proboscia alata* | 0.14 | 53939606 | 76970 |
| MMETSP0816 | Bacillariophyta | *Proboscia inermis* | 0.06 | 51566466 | 28886 |
| MMETSP0327 | Bacillariophyta | *Pseudo-nitzschia delicatissima* | 0.19 | 76463666 | 148478 |
| MMETSP0850 | Bacillariophyta | *Pseudo-nitzschia fraudulenta* | 0.11 | 35381966 | 38254 |
| MMETSP1423 | Bacillariophyta | *Pseudo-nitzschia heimii* | 0.12 | 34362058 | 41362 |
| MMETSP1060 | Bacillariophyta | *Pseudo-nitzschia pungens* | 0.12 | 75981556 | 92662 |
| MMETSP0789 | Bacillariophyta | *Rhizosolenia setigera* | 0.11 | 53486746 | 57924 |
| MMETSP0013_2 | Bacillariophyta | *Skeletonema costatum* | 0.54 | 36792038 | 198180 |
| MMETSP0794_2 | Bacillariophyta | *Stephanopyxis turris* | 0.13 | 28656286 | 38420 |
| MMETSP0800 | Bacillariophyta | *Striatella unipunctata* | 0.13 | 48336382 | 62030 |
| MMETSP0786 | Bacillariophyta | *Thalassionema frauenfeldii* | 0.08 | 51370380 | 40812 |
| MMETSP0156 | Bacillariophyta | *Thalassionema nitzschioides* | 0.12 | 52529482 | 61907 |
| MMETSP0902 | Bacillariophyta | *Thalassiosira antarctica* | 0.35 | 32870602 | 113463 |
| MMETSP0492 | Bacillariophyta | *Thalassiosira gravida* | 0.16 | 39941506 | 65655 |
| MMETSP0737 | Bacillariophyta | *Thalassiosira miniscula* | 0.21 | 72933184 | 149882 |
| MMETSP1067 | Bacillariophyta | *Thalassiosira punctigera* | 0.22 | 56431354 | 124621 |
| MMETSP0403 | Bacillariophyta | *Thalassiosira rotula* | 0.31 | 80967272 | 253484 |
| MMETSP0878 | Bacillariophyta | *Thalassiosira weissflogii* | 0.3 | 52047092 | 154564 |
| MMETSP0152 | Bacillariophyta | *Thalassiothrix antarctica* | 0.1 | 45679326 | 47786 |
| MMETSP1065 | Bacillariophyta | *Amphiprora paludosa* | 0.13 | 66002034 | 87843 |
| MMETSP0169 | Bacillariophyta | *Corethron pennatum* | 0.13 | 51460954 | 65476 |
| MMETSP0010_2 | Bacillariophyta | *Corethron hystrix* | 0.05 | 23183502 | 12379 |
| MMETSP0009_2 | Bacillariophyta | *Grammatophora oceanica* | 0.11 | 31641830 | 34138 |

This file has been converted from its original format for security purposes. Please use CDDD2F8ED7FB9 as a reference.

| | | | | | |
|---|---|---|---|---|---|
| MMETSP0696 | Bacillariophyta | *Extubocellulus spinifer* | 0.14 | 44300740 | 60944 |
| MMETSP1070 | Bacillariophyta | *Minutocellus polymorphus* | 0.13 | 69826230 | 91474 |
| MMETSP1460 | Chlorophyta | *Bathycoccus prasinos* | 0.02 | 55777388 | 10914 |
| MMETSP1392 | Chlorophyta | *Chlamydomonas chlamydogama* | 0.18 | 64172020 | 118213 |
| MMETSP0033_2 | Chlorophyta | *Dolichomastix tenuilepis* | 0.13 | 55347526 | 72640 |
| MMETSP1126 | Chlorophyta | *Dunaliella tertiolecta* | 0.11 | 58248576 | 63375 |
| MMETSP1106 | Chlorophyta | *Mantoniella antarctica* | 0.14 | 48679686 | 66402 |
| MMETSP1401 | Chlorophyta | *Micromonas pusilla* | 0.09 | 43995616 | 37823 |
| MMETSP1327 | Chlorophyta | *Micromonas pusilla* | 0.07 | 45830388 | 30549 |
| MMETSP0034_2 | Chlorophyta | *Nephroselmis pyriformis* | 0.33 | 46610224 | 151760 |
| MMETSP0939 | Chlorophyta | *Ostreococcus lucimarinus* | 0.02 | 62631002 | 13307 |
| MMETSP0929 | Chlorophyta | *Ostreococcus mediterraneus* | 0.04 | 87232616 | 35325 |
| MMETSP0933 | Chlorophyta | *Ostreococcus prasinos* | 0.02 | 54576496 | 13036 |
| MMETSP1161 | Chlorophyta | *Picochlorum oklahomensis* | 0.06 | 41993614 | 27107 |
| MMETSP0941 | Chlorophyta | *Prasinococcus capsulatus* | 0.02 | 47027436 | 10650 |
| MMETSP0806_2 | Chlorophyta | *Prasinoderma coloniale* | 0.03 | 52641318 | 15774 |
| MMETSP1315 | Chlorophyta | *Prasinoderma singularis* | 0.04 | 45156550 | 18933 |
| MMETSP1316 | Chlorophyta | *Pycnococcus provasolii* | 0.03 | 41226934 | 10581 |
| MMETSP0819 | Chlorophyta | *Tetraselmis striata* | 0.09 | 40593864 | 35412 |
| MMETSP0058 | Chlorophyta | *Pyramimonas parkeae* | 0.19 | 67915958 | 130014 |
| MMETSP1050 | Cryptophyta | *Cryptomonas curvata* | 0.07 | 56053624 | 37846 |
| MMETSP0799 | Cryptophyta | *Geminigera cryophila* | 0.05 | 58337084 | 28258 |
| MMETSP1048 | Cryptophyta | *Hanusia phi* | 0.96 | 52458676 | 502153 |
| MMETSP0043_2 | Cryptophyta | *Hemiselmis andersenii* | 0.33 | 78445892 | 262699 |
| MMETSP1356 | Cryptophyta | *Hemiselmis viresens* | 0.08 | 52614122 | 42635 |
| MMETSP1047 | Cryptophyta | *Rhodomonas salina* | 0.11 | 54321504 | 57046 |
| MMETSP0047_2 | Cryptophyta | *Chroomonas mesostigmatica_cf* | 0.06 | 67406484 | 37125 |
| MMETSP1049 | Cryptophyta | *Proteomonas sulcata* | 0.02 | 48515622 | 10005 |
| MMETSP0046_2 | Cryptophyta | *Guillardia theta* | 0.43 | 42158888 | 180684 |
| MMETSP0790 | Dinophyta | *Alexandrium catenella* | 0.01 | 42794264 | 3031 |
| MMETSP0328 | Dinophyta | *Alexandrium minutum* | 0.02 | 22624640 | 5314 |
| MMETSP0378 | Dinophyta | *Alexandrium tamarense* | 0.04 | 28696260 | 11201 |
| MMETSP0258 | Dinophyta | *Amphidinium carterae* | 0.06 | 52247898 | 29626 |
| MMETSP1074 | Dinophyta | *Ceratium fususV* | 0.03 | 54627688 | 18672 |
| MMETSP0797 | Dinophyta | *Dinophysis acuminata* | 0.04 | 48685084 | 20373 |
| MMETSP0116_2 | Dinophyta | *Durinskia baltica* | 0.04 | 46992432 | 20531 |
| MMETSP0503 | Dinophyta | *Heterocapsa rotundata* | 0.13 | 43256526 | 57107 |
| MMETSP0448 | Dinophyta | *Heterocapsa triquestra* | 0.06 | 45359332 | 29357 |
| MMETSP0027 | Dinophyta | *Karenia brevis* | 0.03 | 26748951 | 8238 |
| MMETSP0120_2 | Dinophyta | *Kryptoperidinium foliaceum* | 0.05 | 63629800 | 33936 |
| MMETSP1032 | Dinophyta | *Lingulodinium polyedra* | 0.04 | 65984202 | 24470 |
| MMETSP0370 | Dinophyta | *Peridinium aciculiferum* | 0.03 | 41244260 | 13444 |
| MMETSP0267 | Dinophyta | *Prorocentrum minimum* | 0.06 | 25887799 | 14925 |
| MMETSP0367 | Dinophyta | *Scrippsiella hangoei-like* | 0.04 | 46219246 | 16216 |
| MMETSP0270 | Dinophyta | *Scrippsiella trochoidea* | 0.04 | 36147274 | 16043 |
| MMETSP0229_2 | Dinophyta | *Pyrocystis lunula* | 0 | 62650660 | 1661 |
| MMETSP0796 | Dinophyta | *Pyrodinium bahamense* | 0.03 | 62709420 | 13116 |

This file has been converted from its original format for security purposes. Please use CDDD2F8ED7FB9 as a reference.

| | | | | | |
|---|---|---|---|---|---|
| MMETSP0228 | Dinophyta | *Protoceratium reticulatum* | 0.04 | 46869592 | 17842 |
| MMETSP0784 | Dinophyta | *Gymnodinium catenatum* | 0.05 | 73471400 | 33136 |
| MMETSP0118_2 | Dinophyta | *Glenodinium foliaceum* | 0.07 | 45761730 | 32002 |
| MMETSP0225 | Dinophyta | *Thoracosphaera heimii* | 0 | 54872800 | 1162 |
| MMETSP1036_2 | Dinophyta | *Azadinium spinosum* | 0.15 | 51318154 | 77465 |
| MMETSP1441 | Dinophyta | *Heterocapsa arctica* | 0.09 | 54674410 | 51185 |
| MMETSP1334 | Haptophyta | *Calcidiscus leptoporus* | 0.05 | 44404398 | 20413 |
| MMETSP0143 | Haptophyta | *Chrysochromulina polylepis* | 0.1 | 74214716 | 72977 |
| MMETSP0164_2 | Haptophyta | *Coccolithus pelagicus ssp braarudi* | 0.1 | 53235070 | 52255 |
| MMETSP0994 | Haptophyta | *Emiliania huxleyi* | 0.11 | 25066298 | 28696 |
| MMETSP0944 | Haptophyta | *Isochrysis galbana* | 0.11 | 45534184 | 49496 |
| MMETSP1444 | Haptophyta | *Phaeocystis antarctica* | 0.12 | 47646454 | 59453 |
| MMETSP1465 | Haptophyta | *Phaeocystis cordata* | 0.04 | 38854414 | 15020 |
| MMETSP1136 | Haptophyta | *Pleurochrysis carterae* | 0.11 | 43469752 | 48084 |
| MMETSP1333 | Haptophyta | *Scyphosphaera apsteinii* | 0.03 | 40529618 | 25217 |
| MMETSP1335 | Haptophyta | *Chrysoculter rhomboideus* | 0.07 | 43272560 | 28380 |
| MMETSP0006_2 | Haptophyta | *Prymnesium parvum* | 0.08 | 42932420 | 33478 |
| MMETSP1363 | Haptophyta | *Gephyrocapsa oceanica* | 0.03 | 54668892 | 18456 |
| MMETSP1463 | Haptophyta | *Pavlova lutheri* | 0.05 | 45354612 | 23072 |
| MMETSP1464 | Haptophyta | *Exanthemachrysis gayraliae* | 0.08 | 48832876 | 38546 |
| MMETSP0915 | Ochrophyta | *Aureococcus anophagefferens* | 0.08 | 46827590 | 39555 |
| MMETSP1319 | Ochrophyta | *Bolidomonas pacifica* | 0.18 | 43287000 | 78884 |
| MMETSP0292 | Ochrophyta | *Heterosigma akashiwo* | 0.09 | 32622070 | 28458 |
| MMETSP0888 | Ochrophyta | *Pelagomonas calceolata* | 0.15 | 42273462 | 62406 |
| MMETSP1339 | Ochrophyta | *Fibrocapsa japonica* | 0.11 | 51008030 | 54525 |
| MMETSP1174 | Ochrophyta | *Dictyocha speculum* | 0.07 | 41736862 | 28379 |
| MMETSP1068 | Ochrophyta | *Pseudopedinella elastica* | 0.09 | 59467734 | 52850 |
| MMETSP0947 | Ochrophyta | *Chattonella subsalsa* | 0.04 | 75999072 | 30099 |
| MMETSP1160 | Ochrophyta | *Pinguiococcus pyrenoidosus* | 0.08 | 60971594 | 46308 |
| MMETSP1323 | Ochrophyta | *Florenciella parvula* | 0.28 | 24016606 | 66145 |
| MMETSP1163 | Ochrophyta | *Phaeomonas parva* | 0.15 | 47821364 | 69,830 |
| MMETSP0890 | Ochrophyta | *Aureoumbra lagunensis* | 0.05 | 24957712 | 12204 |
| MMETSP0882 | Ochrophyta | *Pelagococcus subviridis* | 0.07 | 36185352 | 26542 |
| MMETSP0167 | Rhodophyta | *Rhodella maculata* | 0.03 | 54377336 | 17657 |
| MMETSP0011_2 | Rhodophyta | *Rhodosorus marinus* | 0.08 | 42420550 | 35037 |

This file has been converted from its original format for security purposes. Please use CDDD2F8ED7FB9 as a reference.
