## Supplementary material for "The potential role of viruses controlling phytoplankton community size structure": List of 125 phytoplankton species with their MMETSP identifiers and cell volumes along with the references.

Table I: List of 125 phytoplankton species with their MMETSP identifiers and cell volumes along with the references.

| No. | MMETSP ID | Phylum | Species | Cell volume ($\mu m^3$ $cell^{-1}$) | Reference |
|---|---|---|---|---|---|
| 1 | MMETSP0316 | Bacillariophyta | *Amphora coffeaeformis* | 2826.00 | Olenina, I. et al. Biovolumes and size-classes of phytoplankton in the Baltic Sea HELCOM Balt.Sea Environ. Proc. 106 (2006). |
| 2 | MMETSP0705 | Bacillariophyta | *Asterionellopsis glacialis* | 1381.07 | Sal, S., López-Urrutia, A., Irigoien, X., Harbour, D. S. & Harris, R. P. Marine microplankton diversity database. *Ecology* 94, doi:http://dx.doi.org/10.1890/13-0236.1 (2013). |
| 3 | MMETSP1449 | Bacillariophyta | *Attheya septentrionalis* | 100.00 | Olenina, I. et al. Biovolumes and size-classes of phytoplankton in the Baltic Sea HELCOM Balt.Sea Environ. Proc. 106 (2006). |
| 4 | MMETSP1435 | Bacillariophyta | *Chaetoceros brevis* | 4227.60 | Olenina, I. et al. Biovolumes and size-classes of phytoplankton in the Baltic Sea HELCOM Balt.Sea Environ. Proc. 106 (2006). |
| 5 | MMETSP1336 | Bacillariophyta | *Chaetoceros cf. neogracile* | 179.59 | Waite, A., Gallager, S. & Dam, H. G. New measurements of phytoplankton aggregation in a flocculator using videography and image analysis. *Marine Ecology Progress Series* 155, 77-88, doi:Doi 10.3354/Meps155077 (1997). |
| 6 | MMETSP0150 | Bacillariophyta | *Chaetoceros debilis* | 2791.40 | Olenina, I. et al. Biovolumes and size-classes of phytoplankton in the Baltic Sea HELCOM Balt.Sea Environ. Proc. 106 (2006). |
| 7 | MMETSP1447 | Bacillariophyta | *Chaetoceros dichaeta* | 609.80 | Sal, S., López-Urrutia, A., Irigoien, X., Harbour, D. S. & Harris, R. P. Marine microplankton diversity database. *Ecology* 94, doi:http://dx.doi.org/10.1890/13-0236.1 (2013). |
| 8 | MMETSP1066 | Bacillariophyta | *Coscinodiscus wailesii* | 9277758.00 | Olenina, I. et al. Biovolumes and size-classes of phytoplankton in the Baltic Sea HELCOM Balt.Sea Environ. Proc. 106 (2006). |
| 9 | MMETSP1057 | Bacillariophyta | *Cyclotella meneghiniana* | 10161.00 | Olenina, I. et al. Biovolumes and size-classes of phytoplankton in the Baltic Sea HELCOM Balt.Sea Environ. Proc. 106 (2006). |
| 10 | MMETSP0017_2 | Bacillariophyta | *Cylindrotheca closterium* | 254.00 | Olenina, I. et al. Biovolumes and size-classes of phytoplankton in the Baltic Sea HELCOM Balt.Sea Environ. Proc. 106 (2006). |
| 11 | MMETSP0580 | Bacillariophyta | *Dactyliosolen fragilissimus* | 8658.20 | Olenina, I. et al. Biovolumes and size-classes of phytoplankton in the Baltic Sea HELCOM Balt.Sea Environ. Proc. 106 (2006). |
| 12 | MMETSP1058 | Bacillariophyta | *Detonula confervacea* | 1606.33 | Olenina, I. et al. Biovolumes and size-classes of phytoplankton in the Baltic Sea HELCOM Balt.Sea Environ. Proc. 106 (2006). |
| 13 | MMETSP0998 | Bacillariophyta | *Ditylum brightwellii* | 46603.25 | Olenina, I. et al. Biovolumes and size-classes of phytoplankton in the Baltic Sea HELCOM Balt.Sea Environ. Proc. 106 (2006). |
| 14 | MMETSP1437 | Bacillariophyta | *Eucampia antarctica* | 64014.00 | Kang, S. H. *et al.* Antarctic phytoplankton assemblages in the marginal ice zone of the northwestern Weddell Sea. *J Plankton Res* **23**, 333-352, doi:DOI 10.1093/plankt/23.4.333 (2001). |

This file has been converted from its original format for security purposes. Please use C56CD658F48A1 as a reference.

| | | | | | |
|---|---|---|---|---|---|
| 15 | MMETSP0733 | Bacillariophyta | *Fragilariopsis kerguelensis* | 603.30 | Sal, S., López-Urrutia, A., Irigoien, X., Harbour, D. S. & Harris, R. P. Marine microplankton diversity database. Ecology 94, doi:http://dx.doi.org/10.1890/13-0236.1 (2013). |
| 16 | MMETSP1171 | Bacillariophyta | *Helicotheca tamensis* | 6007.88 | Sal, S., López-Urrutia, A., Irigoien, X., Harbour, D. S. & Harris, R. P. Marine microplankton diversity database. Ecology 94, doi:http://dx.doi.org/10.1890/13-0236.1 (2013). |
| 17 | MMETSP1362 | Bacillariophyta | *Leptocylindrus danicus* | 1338.78 | Olenina, I. et al. Biovolumes and size-classes of phytoplankton in the Baltic Sea HELCOM Balt.Sea Environ. Proc. 106 (2006). |
| 18 | MMETSP0744 | Bacillariophyta | *Nitzschia punctata* | 7713.00 | http://diatom.ansp.org/taxaservice/ShowTaxon.ashx?naded_id=185039 |
| 19 | MMETSP0015_2 | Bacillariophyta | *Odontella aurita* | 21760.00 | Olenina, I. et al. Biovolumes and size-classes of phytoplankton in the Baltic Sea HELCOM Balt.Sea Environ. Proc. 106 (2006). |
| 20 | MMETSP0160_2 | Bacillariophyta | *Odontella Sinensis* | 1000875.00 | Olenina, I. et al. Biovolumes and size-classes of phytoplankton in the Baltic Sea HELCOM Balt.Sea Environ. Proc. 106 (2006). |
| 21 | MMETSP0174 | Bacillariophyta | *Proboscia alata* | 62530.00 | Sal, S., López-Urrutia, A., Irigoien, X., Harbour, D. S. & Harris, R. P. Marine microplankton diversity database. *Ecology* 94, doi:http://dx.doi.org/10.1890/13-0236.1 (2013). |
| 22 | MMETSP0816 | Bacillariophyta | *Proboscia inermis* | 12566.00 | Sal, S., López-Urrutia, A., Irigoien, X., Harbour, D. S. & Harris, R. P. Marine microplankton diversity database. *Ecology* 94, doi:http://dx.doi.org/10.1890/13-0236.1 (2013). |
| 23 | MMETSP0327 | Bacillariophyta | *Pseudo-nitzschia delicatissima* | 168.33 | Olenina, I. et al. Biovolumes and size-classes of phytoplankton in the Baltic Sea HELCOM Balt.Sea Environ. Proc. 106 (2006). |
| 24 | MMETSP0850 | Bacillariophyta | *Pseudo-nitzschia fraudulenta* | 149.64 | Sal, S., López-Urrutia, A., Irigoien, X., Harbour, D. S. & Harris, R. P. Marine microplankton diversity database. *Ecology* 94, doi:http://dx.doi.org/10.1890/13-0236.1 (2013). |
| 25 | MMETSP1423 | Bacillariophyta | *Pseudo-nitzschia heimii* | 965.00 | Kang, S. H. *et al.* Antarctic phytoplankton assemblages in the marginal ice zone of the northwestern Weddell Sea. *J Plankton Res* **23**, 333-352, doi:DOI 10.1093/plankt/23.4.333 (2001). |
| 26 | MMETSP1060 | Bacillariophyta | *Pseudo-nitzschia pungens* | 2266.00 | Olenina, I. et al. Biovolumes and size-classes of phytoplankton in the Baltic Sea HELCOM Balt.Sea Environ. Proc. 106 (2006). |
| 27 | MMETSP0789 | Bacillariophyta | *Rhizosolenia setigera* | 31600.00 | Sal, S., López-Urrutia, A., Irigoien, X., Harbour, D. S. & Harris, R. P. Marine microplankton diversity database. *Ecology* 94, doi:http://dx.doi.org/10.1890/13-0236.1 (2013). |
| 28 | MMETSP0013_2 | Bacillariophyta | *Skeletonema costatum* | 388.80 | Olenina, I. et al. Biovolumes and size-classes of phytoplankton in the Baltic Sea HELCOM Balt.Sea Environ. Proc. 106 (2006). |
| 29 | MMETSP0794_2 | Bacillariophyta | *Stephanopyxis turris* | 28260.00 | Olenina, I. et al. Biovolumes and size-classes of phytoplankton in the Baltic Sea HELCOM Balt.Sea Environ. Proc. 106 (2006). |
| 30 | MMETSP0800 | Bacillariophyta | *Striatella unipunctata* | 4339.05 | Sal, S., López-Urrutia, A., Irigoien, X., Harbour, D. S. & Harris, R. P. Marine microplankton diversity database. *Ecology* 94, doi:http://dx.doi.org/10.1890/13-0236.1 (2013). |

This file has been converted from its original format for security purposes. Please use C56CD658F48A1 as a reference.

| | | | | | |
|---|---|---|---|---|---|
| 31 | MMETSP0786 | Bacillariophyta | *Thalassionema frauenfeldii* | 480.00 | Sal, S., López-Urrutia, A., Irigoien, X., Harbour, D. S. & Harris, R. P. Marine microplankton diversity database. *Ecology* 94, doi:http://dx.doi.org/10.1890/13-0236.1 (2013). |
| 32 | MMETSP0156 | Bacillariophyta | *Thalassionema nitzschioides* | 962.13 | Olenina, I. et al. Biovolumes and size-classes of phytoplankton in the Baltic Sea HELCOM Balt.Sea Environ. Proc. 106 (2006). |
| 33 | MMETSP0902 | Bacillariophyta | *Thalassiosira antarctica* | 785.40 | Sal, S., López-Urrutia, A., Irigoien, X., Harbour, D. S. & Harris, R. P. Marine microplankton diversity database. *Ecology* 94, doi:http://dx.doi.org/10.1890/13-0236.1 (2013). |
| 34 | MMETSP0492 | Bacillariophyta | *Thalassiosira gravida* | 16632.00 | Olenina, I. et al. Biovolumes and size-classes of phytoplankton in the Baltic Sea HELCOM Balt.Sea Environ. Proc. 106 (2006). |
| 35 | MMETSP0737 | Bacillariophyta | *Thalassiosira miniscula* | 1145.00 | von Dassow, P., Petersen, T. W., Chepurnov, V. A. & Armbrust, E. V. Inter- and intraspecific relationships between nuclear DNA content and cell size in selected members of the centric diatom genus Thalassiosira (Bacillariophyceae). *J Phycol* 44, 335-349, doi:DOI 10.1111/j.1529-8817.2008.00476.x (2008). |
| 36 | MMETSP1067 | Bacillariophyta | *Thalassiosira punctigera* | 69619.67 | Olenina, I. et al. Biovolumes and size-classes of phytoplankton in the Baltic Sea HELCOM Balt.Sea Environ. Proc. 106 (2006). |
| 37 | MMETSP0403 | Bacillariophyta | *Thalassiosira rotula* | 15072.00 | Olenina, I. et al. Biovolumes and size-classes of phytoplankton in the Baltic Sea HELCOM Balt.Sea Environ. Proc. 106 (2006). |
| 38 | MMETSP0878 | Bacillariophyta | *Thalassiosira weissflogii* | 2950.00 | Olenina, I. et al. Biovolumes and size-classes of phytoplankton in the Baltic Sea HELCOM Balt.Sea Environ. Proc. 106 (2006). |
| 39 | MMETSP0152 | Bacillariophyta | *Thalassiothrix antarctica* | 7500.00 | Sal, S., López-Urrutia, A., Irigoien, X., Harbour, D. S. & Harris, R. P. Marine microplankton diversity database. *Ecology* 94, doi:http://dx.doi.org/10.1890/13-0236.1 (2013). |
| 40 | MMETSP1065 | Bacillariophyta | *Amphiprora paludosa* | 40857.00 | Andreoli, C. *et al.* Diatoms and Dinoflagellates in Terra-Nova Bay (Ross Sea Antarctica) during Austral Summer 1990. *Polar Biology* **15**, 465-475 (1995). |
| 41 | MMETSP0169 | Bacillariophyta | *Corethron pennatum* | 38000.00 | Timmermans, K. R., van der Wagt, B. & de Baar, H. J. W. Growth rates, half-saturation constants, and silicate, nitrate, and phosphate depletion in relation to iron availability of four large, open-ocean diatoms from the Southern Ocean. *Limnol Oceanogr* **49**, 2141-2151 (2004). |
| 42 | MMETSP0010_2 | Bacillariophyta | *Corethron hystrix* | 6492.00 | Hitchcock, G. L. An Examination of Diatom Area - Volume Ratios and Their Influence on Estimates of Plasma-Volume. *J Plankton Res* **5**, 311-324, doi:DOI 10.1093/plankt/5.3.311 (1983). |
| 43 | MMETSP0009_2 | Bacillariophyta | *Grammatophora oceanica* | 53.01 | Martin, C. L. & Tortell, P. D. Bicarbonate transport and extracellular carbonic anhydrase in marine diatoms. *Physiol Plantarum* **133**, 106-116, doi:DOI 10.1111/j.1399-3054.2008.01054.x (2008). |
| 44 | MMETSP0696 | Bacillariophyta | *Extubocellulus spinifer* | 31.91 | http://nordicmicroalgae.org/taxon/Extubocellulus%20spinifer |
| 45 | MMETSP1070 | Bacillariophyta | *Minutocellus polymorphus* | 16.10 | Sarno, D., Zingone, A., Saggiomo, V. & Carrada, G. C. Phytoplankton Biomass and Species Composition in a Mediterranean Coastal Lagoon. *Hydrobiologia* **271**, 27-40, doi:Doi 10.1007/Bf00005692 (1993). |
| 46 | MMETSP1460 | Chlorophyta | *Bathycoccus prasinos* | 2.36 | Eikrem, W. & Throndsen, J. The Ultrastructure of Bathycoccus Gen-Nov and Bathycoccus-Prasinos Sp-Nov, a Nonmotile Picoplanktonic Alga (Chlorophyta, Prasinophyceae) from the Mediterranean and Atlantic. *Phycologia* **29**, 344-350, doi:DOI |

This file has been converted from its original format for security purposes. Please use C56CD658F48A1 as a reference.

| | | | | | |
|---|---|---|---|---|---|
| | | | | | 10.2216/i0031-8884-29-3-344.1 (1990). |
| 47 | MMETSP1392 | Chlorophyta | *Chlamydomonas chlamydogama* | 508.00 | Bold, H. C. The morphology of Chlamydomonas chlamydogama, sp. nov. *Bulletin of the Torrey Botanical Club*, 101-108 (1949). |
| 48 | MMETSP0033_2 | Chlorophyta | *Dolichomastix tenuilepis* | 28.00 | Throndsen, J. & Zingone, A. Dolichomastix tenuilepis sp nov., a first insight into the microanatomy of the genus Dolichomastix (Mamiellales, Prasinophyceae, Chlorophyta). *Phycologia* **36**, 244-254, doi:DOI 10.2216/i0031-8884-36-3-244.1 (1997). |
| 49 | MMETSP1126 | Chlorophyta | *Dunaliella tertiolecta* | 212.00 | Meyer, B., Irigoien, X., Graeve, M., Head, R. N. & Harris, R. P. Feeding rates and selectivity among nauplii, copepodites and adult females of Calanus finmarchicus and Calanus helgolandicus. *Helgoland Mar Res* **56**, 169-176, doi:DOI 10.1007/s10152-002-0105-3 (2002). |
| 50 | MMETSP1106 | Chlorophyta | *Mantoniella antarctica* | 29.00 | Marchant, H. J., Buck, K. R., Garrison, D. L. & Thomsen, H. A. Mantoniella in Antarctic Waters Including the Description of Mantoniella-Antarctica Sp-Nov (Prasinophyceae). *J Phycol* **25**, 167-174, doi:DOI 10.1111/j.0022-3646.1989.00167.x (1989). |
| 51 | MMETSP1401 | Chlorophyta | *Micromonas pusilla CCAC1681* | 1.00 | Olenina, I. et al. Biovolumes and size-classes of phytoplankton in the Baltic Sea HELCOM Balt.Sea Environ. Proc. 106 (2006). |
| 52 | MMETSP1327 | Chlorophyta | *Micromonas pusilla RCC2306* | 1.00 | Olenina, I. et al. Biovolumes and size-classes of phytoplankton in the Baltic Sea HELCOM Balt.Sea Environ. Proc. 106 (2006). |
| 53 | MMETSP0034_2 | Chlorophyta | *Nephroselmis pyriformis* | 56.00 | Gismervik, I. Top-down impact by copepods on ciliate numbers and persistence depends on copepod and ciliate species composition. *J Plankton Res* **28**, 499-507, doi:DOI 10.1093/plankt/fbi135 (2006). |
| 54 | MMETSP0939 | Chlorophyta | *Ostreococcus lucimarinus* | 0.60 | http://roscoff-culture-collection.org/ |
| 55 | MMETSP0929 | Chlorophyta | *Ostreococcus mediterraneus* | 1.07 | Subirana, L. *et al.* Morphology, Genome Plasticity, and Phylogeny in the Genus Ostreococcus Reveal a Cryptic Species, O. mediterraneus sp nov (Mamiellales, Mamiellophyceae). *Protist* **164**, 643-659, doi:DOI 10.1016/j.protis.2013.06.002 (2013). |
| 56 | MMETSP0933 | Chlorophyta | *Ostreococcus prasinos* | 1.00 | http://roscoff-culture-collection.org/ |
| 57 | MMETSP1161 | Chlorophyta | *Picochlorum oklahomensis* | 4.20 | Zhu, Y. & Dunford, N. T. Growth and Biomass Characteristics of Picochlorum oklahomensis and Nannochloropsis oculata. *J Am Oil Chem Soc* **90**, 841-849, doi:DOI 10.1007/s11746-013-2225-0 (2013). |
| 58 | MMETSP0941 | Chlorophyta | *Prasinococcus capsulatus* | 33.50 | http://roscoff-culture-collection.org/ |
| 59 | MMETSP0806_2 | Chlorophyta | *Prasinoderma coloniale* | 14.00 | http://roscoff-culture-collection.org/ |
| 60 | MMETSP1315 | Chlorophyta | *Prasinoderma singularis* | 14.00 | http://roscoff-culture-collection.org/ |
| 61 | MMETSP1316 | Chlorophyta | *Pycnococcus provasolii* | 14.00 | http://roscoff-culture-collection.org/ |
| 62 | MMETSP0819 | Chlorophyta | *Tetraselmis striata* | 382.00 | http://roscoff-culture-collection.org/ |

This file has been converted from its original format for security purposes. Please use C56CD658F48A1 as a reference.

| | | | | | |
|---|---|---|---|---|---|
| 63 | MMETSP0058 | Chlorophyta | *Pyramimonas parkeae* | 600.00 | Kohata, K. & Watanabe, M. Diel Changes in the Composition of Photosynthetic Pigments and Cellular Carbon and Nitrogen in Pyraminonas-Parkeae (Prasinophyceae). *J Phycol* **25**, 377-385, doi:DOI 10.1111/j.1529-8817.1989.tb00134.x (1989). |
| 64 | MMETSP1050 | Cryptophyta | *Cryptomonas curvata* | 11751.00 | Rocha, O. & Duncan, A. The Relationship between Cell Carbon and Cell-Volume in Fresh-Water Algal Species Used in Zooplanktonic Studies. *J Plankton Res* **7**, 279-294, doi:DOI 10.1093/plankt/7.2.279 (1985). |
| 65 | MMETSP0799 | Cryptophyta | *Geminigera cryophila* | 301.59 | Hill, D. R. A. A Revised Circumscription of Cryptomonas (Cryptophyceae) Based on Examination of Australian Strains. *Phycologia* **30**, 170-188, doi:DOI 10.2216/i0031-8884-30-2-170.1 (1991). |
| 66 | MMETSP1048 | Cryptophyta | *Hanusia phi* | 62.83 | Deane, J. A., Hill, D. R. A., Brett, S. J. & McFadden, G. I. Hanusia phi gen. et sp. nov. (Cryptophyceae): characterization of 'Cryptomonas sp. Phi'. *Eur J Phycol* **33**, 149-154, doi:Doi 10.1017/S0967026298001553 (1998). |
| 67 | MMETSP0043_2 | Cryptophyta | *Hemiselmis andersenii* | 87.96 | Lane, C. E. & Archibald, J. M. New marine members of the genus Hemiselmis (Cryptomonadales, Cryptophyceae). *J Phycol* **44**, 439-450, doi:DOI 10.1111/j.1529-8817.2008.00486.x (2008). |
| 68 | MMETSP1356 | Cryptophyta | *Hemiselmis viresens* | 15.00 | Olenina, I. et al. Biovolumes and size-classes of phytoplankton in the Baltic Sea HELCOM Balt.Sea Environ. Proc. 106 (2006). |
| 69 | MMETSP1047 | Cryptophyta | *Rhodomonas salina* | 141.00 | Olenina, I. et al. Biovolumes and size-classes of phytoplankton in the Baltic Sea HELCOM Balt.Sea Environ. Proc. 106 (2006). |
| 70 | MMETSP0047_2 | Cryptophyta | *Chroomonas mesostigmatica_cf* | 130.90 | https://ncma.bigelow.org/ccmp1168 |
| 71 | MMETSP1049 | Cryptophyta | *Proteomonas sulcata* | 130.90 | Hill, D. R. A. & Wetherbee, R. Proteomonas-Sulcata Gen Et Sp-Nov (Cryptophyceae), a Cryptomonad with 2 Morphologically Distinct and Alternating Forms. *Phycologia* **25**, 521-543, doi:DOI 10.2216/i0031-8884-25-4-521.1 (1986). |
| 72 | MMETSP0046_2 | Cryptophyta | *Guillardia theta* | 23.56 | Deane, J. A., Hill, D. R. A., Brett, S. J. & McFadden, G. I. Hanusia phi gen. et sp. nov. (Cryptophyceae): characterization of 'Cryptomonas sp. Phi'. *Eur J Phycol* **33**, 149-154, doi:Doi 10.1017/S0967026298001553 (1998). |
| 73 | MMETSP0790 | Dinophyta | *Alexandrium catenella* | 13027.00 | Menden-Deuer, S. & Lessard, E. J. Carbon to volume relationships for dinoflagellates, diatoms, and other protist plankton. *Limnol Oceanogr* **45**, 569-579 (2000). |
| 74 | MMETSP0328 | Dinophyta | *Alexandrium minutum* | 9156.50 | Olenina, I. et al. Biovolumes and size-classes of phytoplankton in the Baltic Sea HELCOM Balt.Sea Environ. Proc. 106 (2006). |
| 75 | MMETSP0378 | Dinophyta | *Alexandrium tamarense* | 14337.67 | Olenina, I. et al. Biovolumes and size-classes of phytoplankton in the Baltic Sea HELCOM Balt.Sea Environ. Proc. 106 (2006). |
| 76 | MMETSP0258 | Dinophyta | *Amphidinium carterae* | 432.00 | Olenina, I. et al. Biovolumes and size-classes of phytoplankton in the Baltic Sea HELCOM Balt.Sea Environ. Proc. 106 (2006). |
| 77 | MMETSP1074 | Dinophyta | *Ceratium fususV* | 14500.00 | Olenina, I. et al. Biovolumes and size-classes of phytoplankton in the Baltic Sea HELCOM Balt.Sea Environ. Proc. 106 (2006). |
| 78 | MMETSP0797 | Dinophyta | *Dinophysis acuminata* | 14194.00 | Sal, S., López-Urrutia, A., Irigoien, X., Harbour, D. S. & Harris, R. P. Marine microplankton diversity database. *Ecology* 94, doi:http://dx.doi.org/10.1890/13-0236.1 (2013). |

This file has been converted from its original format for security purposes. Please use C56CD658F48A1 as a reference.

| | | | | | |
|---|---|---|---|---|---|
| 79 | MMETSP0116_2 | Dinophyta | *Durinskia baltica* | 7065.00 | Olenina, I. et al. Biovolumes and size-classes of phytoplankton in the Baltic Sea HELCOM Balt.Sea Environ. Proc. 106 (2006). |
| 80 | MMETSP0503 | Dinophyta | *Heterocapsa rotundata* | 234.00 | Olenina, I. et al. Biovolumes and size-classes of phytoplankton in the Baltic Sea HELCOM Balt.Sea Environ. Proc. 106 (2006). |
| 81 | MMETSP0448 | Dinophyta | *Heterocapsa triquestra* | 1670.17 | Olenina, I. et al. Biovolumes and size-classes of phytoplankton in the Baltic Sea HELCOM Balt.Sea Environ. Proc. 106 (2006). |
| 82 | MMETSP0027 | Dinophyta | *Karenia brevis* | 6494.28 | http://www.sms.si.edu/irlspec/Kareni_brevis.htm |
| 83 | MMETSP0120_2 | Dinophyta | *Kryptoperidinium foliaceum* | 10598.00 | Olenina, I. et al. Biovolumes and size-classes of phytoplankton in the Baltic Sea HELCOM Balt.Sea Environ. Proc. 106 (2006). |
| 84 | MMETSP1032 | Dinophyta | *Lingulodinium polyedra* | 24982.25 | Olenina, I. et al. Biovolumes and size-classes of phytoplankton in the Baltic Sea HELCOM Balt.Sea Environ. Proc. 106 (2006). |
| 85 | MMETSP0370 | Dinophyta | *Peridinium aciculiferum* | 8500.00 | Rengefors, K. & Legrand, C. Broad allelopathic activity in Peridinium aciculiferum (Dinophyceae). *Eur J Phycol* **42**, 341-349, doi:Doi 10.1080/09670260701529604 (2007). |
| 86 | MMETSP0267 | Dinophyta | *Prorocentrum minimum* | 1240.33 | Olenina, I. et al. Biovolumes and size-classes of phytoplankton in the Baltic Sea HELCOM Balt.Sea Environ. Proc. 106 (2006). |
| 87 | MMETSP0367 | Dinophyta | *Scrippsiella hangoei-like* | 6874.50 | Olenina, I. et al. Biovolumes and size-classes of phytoplankton in the Baltic Sea HELCOM Balt.Sea Environ. Proc. 106 (2006). |
| 88 | MMETSP0270 | Dinophyta | *Scrippsiella trochoidea* | 4408.00 | Menden-Deuer, S., Lessard, E. J. & Satterberg, J. Effect of preservation on dinoflagellate and diatom cell volume and consequences for carbon biomass predictions. *Marine Ecology Progress Series* **222**, 41-50, doi:Doi 10.3354/Meps222041 (2001). |
| 89 | MMETSP0229_2 | Dinophyta | *Pyrocystis lunula* | 118984.38 | Swift, E. & Meunier, V. Effects of Light-Intensity on Division Rate, Stimulable Bioluminescence and Cell-Size of Oceanic Dinoflagellates Dissodinium-Lunula, Pyrocystis-Fusiformis and Pyrocystis-Noctiluca. *J Phycol* **12**, 14-22, doi:DOI 10.1111/j.0022-3646.1976.00014.x (1976). |
| 90 | MMETSP0796 | Dinophyta | *Pyrodinium bahamense* | 35427.00 | Steidinger, K. A., Tester, L. S. & Taylor, F. J. R. A Redescription of Pyrodinium-Bahamense Var Compressa (Bohm) Stat Nov from Pacific Red Tides. *Phycologia* **19**, 329-334, doi:DOI 10.2216/i0031-8884-19-4-329.1 (1980). |
| 91 | MMETSP0228 | Dinophyta | *Protoceratium reticulatum* | 16983.25 | http://nordicmicroalgae.org/taxon/Protoceratium%20reticulatum |
| 92 | MMETSP0784 | Dinophyta | *Gymnodinium catenatum* | 14744.58 | Moreygaines, G. Gymnodinium-Catenatum Graham (Dinophyceae) - Morphology and Affinities with Armored Forms. *Phycologia* **21**, 154-163, doi:DOI 10.2216/i0031-8884-21-2-154.1 (1982). |
| 93 | MMETSP1036_2 | Dinophyta | *Azadinium spinosum* | 660.93 | Tillmann, U., Elbrachter, M., Krock, B., John, U. & Cembella, A. Azadinium spinosum gen. et sp nov (Dinophyceae) identified as a primary producer of azaspiracid toxins. *Eur J Phycol* **44**, 63-79, doi:Pii 909484059 Doi 10.1080/09670260802578534 (2009). |

This file has been converted from its original format for security purposes. Please use C56CD658F48A1 as a reference.

| | | | | | |
|---|---|---|---|---|---|
| 94 | MMETSP0118_2 | Dinophyta | *Glenodinium foliaceum* | 8163.00 | Menden-Deuer, S. & Lessard, E. J. Carbon to volume relationships for dinoflagellates, diatoms, and other protist plankton. *Limnol Oceanogr* **45**, 569-579 (2000). |
| 95 | MMETSP0225 | Dinophyta | *Thoracosphaera heimii* | 1353.00 | Ho, T. Y. *et al.* The elemental composition of some marine phytoplankton. *J Phycol* **39**, 1145-1159, doi:DOI 10.1111/j.0022-3646.2003.03-090.x (2003). |
| 96 | MMETSP1441 | Dinophyta | *Heterocapsa arctica* | 1042.74 | Horiguchi, T. Heterocapsa arctica sp nov (Peridiniales, Dinophyceae), a new marine dinoflagellate from the arctic. *Phycologia* **36**, 488-491, doi:DOI 10.2216/i0031-8884-36-6-488.1 (1997). |
| 97 | MMETSP1334 | Haptophyta | *Calcidiscus leptoporus* | 1060.00 | Sal, S., López-Urrutia, A., Irigoien, X., Harbour, D. S. & Harris, R. P. Marine microplankton diversity database. *Ecology* 94, doi:http://dx.doi.org/10.1890/13-0236.1 (2013). |
| 98 | MMETSP0143 | Haptophyta | *Chrysochromulina polylepis* | 271.67 | Olenina, I. et al. Biovolumes and size-classes of phytoplankton in the Baltic Sea HELCOM Balt.Sea Environ. Proc. 106 (2006). |
| 99 | MMETSP0164_2 | Haptophyta | *Coccolithus pelagicus ssp braarudi* | 192.50 | Sal, S., López-Urrutia, A., Irigoien, X., Harbour, D. S. & Harris, R. P. Marine microplankton diversity database. *Ecology* 94, doi:http://dx.doi.org/10.1890/13-0236.1 (2013). |
| 100 | MMETSP0994 | Haptophyta | *Emiliania huxleyi* | 39.50 | Olenina, I. et al. Biovolumes and size-classes of phytoplankton in the Baltic Sea HELCOM Balt.Sea Environ. Proc. 106 (2006). |
| 101 | MMETSP0944 | Haptophyta | *Isochrysis galbana* | 47.00 | Flynn, K. J., Davidson, K. & Cunningham, A. Prey selection and rejection by a microflagellate; Implications for the study and operation of microbial food webs. *J Exp Mar Biol Ecol* **196**, 357-372, doi:Doi 10.1016/0022-0981(95)00140-9(1996). |
| 102 | MMETSP1444 | Haptophyta | *Phaeocystis antarctica* | 19.00 | Kang, S. H. *et al.* Antarctic phytoplankton assemblages in the marginal ice zone of the northwestern Weddell Sea. *J Plankton Res* **23**, 333-352, doi:DOI 10.1093/plankt/23.4.333 (2001). |
| 103 | MMETSP1465 | Haptophyta | *Phaeocystis cordata* | 14.00 | Sal, S., López-Urrutia, A., Irigoien, X., Harbour, D. S. & Harris, R. P. Marine microplankton diversity database. *Ecology* 94, doi:http://dx.doi.org/10.1890/13-0236.1 (2013). |
| 104 | MMETSP1136 | Haptophyta | *Pleurochrysis carterae* | 735.00 | Olenina, I. et al. Biovolumes and size-classes of phytoplankton in the Baltic Sea HELCOM Balt.Sea Environ. Proc. 106 (2006). |
| 105 | MMETSP1333 | Haptophyta | *Scyphosphaera apsteinii* | 1767.15 | http://roscoff-culture-collection.org/rcc-strain-details/1480 |
| 106 | MMETSP1335 | Haptophyta | *Chrysoculter rhomboideus* | 206.76 | Nakayama, T., Yoshida, M., Noel, M. H., Kawachi, M. & Inouye, I. Ultrastructure and phylogenetic position of Chrysoculter rhomboideus gen. et sp nov (Prymnesiophyceae), a new flagellate haptophyte from Japanese coastal waters. *Phycologia* **44**, 369-383, doi:Doi 10.2216/0031-8884(2005)44[369:Uappoc]2.0.Co;2 (2005). |
| 107 | MMETSP0006_2 | Haptophyta | *Prymnesium parvum* | 170.50 | Blossom, H. E. e. a. Prymnesium parvum revisited: Relationship between allelopathy, ichthyotoxicity, and chemical profiles in 5 strains. *Aquatic toxicology* **157**, 159-166 (2014). |
| 108 | MMETSP1363 | Haptophyta | *Gephyrocapsa oceanica* | 142.00 | Ho, T. Y. *et al.* The elemental composition of some marine phytoplankton. *J Phycol* **39**, 1145-1159, doi:DOI 10.1111/j.0022-3646.2003.03-090.x (2003). |
| 109 | MMETSP1463 | Haptophyta | *Pavlova lutheri* | 91.00 | Volkman, J. K., Jeffrey, S. W., Nichols, P. D., Rogers, G. I. & Garland, C. D. Fatty-Acid and Lipid-Composition of 10 Species |

This file has been converted from its original format for security purposes. Please use C56CD658F48A1 as a reference.

| | | | | | |
|---|---|---|---|---|---|
| | | | | | of Microalgae Used in Mariculture. *J Exp Mar Biol Ecol* **128**, 219-240, doi:Doi 10.1016/0022-0981(89)90029-4 (1989). |
| 110 | MMETSP1464 | Haptophyta | *Exanthemachrysis gayraliae* | 65.45 | http://roscoff-culture-collection.org/ |
| 111 | MMETSP0915 | Ochrophyta | *Aureococcus anophagefferens* | 4.20 | Bricelj, V. M. & Lonsdale, D. J. Aureococcus anophagefferens: Causes and ecological consequences of brown tides in US mid-Atlantic coastal waters. *Limnol Oceanogr* **42**, 1023-1038 (1997). |
| 112 | MMETSP1319 | Ochrophyta | *Bolidomonas pacifica* | 0.90 | Guillou, L. *et al.* Bolidomonas: A new genus with two species belonging to a new algal class, the Bolidophyceae (Heterokonta). *J Phycol* **35**, 368-381, doi:DOI 10.1046/j.1529-8817.1999.3520368.x (1999). |
| 113 | MMETSP0292 | Ochrophyta | *Heterosigma akashiwo* | 4187.00 | Olenina, I. et al. Biovolumes and size-classes of phytoplankton in the Baltic Sea HELCOM Balt.Sea Environ. Proc. 106 (2006). |
| 114 | MMETSP0888 | Ochrophyta | *Pelagomonas calceolata* | 3.53 | Andersen, R. A., Saunders, G. W., Paskind, M. P. & Sexton, J. P. Ultrastructure and 18s Ribosomal-Rna Gene Sequence for Pelagomonas-Calceolata Gen Et Sp-Nov and the Description of a New Algal Class, the Pelagophyceae Classis Nov. *J Phycol* **29**, 701-715, doi:DOI 10.1111/j.0022-3646.1993.00701.x (1993). |
| 115 | MMETSP1339 | Ochrophyta | *Fibrocapsa japonica* | 4400.00 | de Boer, M. K., Tyl, M. R., Vrieling, E. G. & van Rijssel, M. Effects of salinity and nutrient conditions on growth and haemolytic activity of Fibrocapsa japonica (Raphidophyceae). *Aquat Microb Ecol* **37**, 171-181, doi:Doi 10.3354/Ame037171 (2004). |
| 116 | MMETSP1174 | Ochrophyta | *Dictyocha speculum* | 2093.00 | Olenina, I. et al. Biovolumes and size-classes of phytoplankton in the Baltic Sea HELCOM Balt.Sea Environ. Proc. 106 (2006). |
| 117 | MMETSP1068 | Ochrophyta | *Pseudopedinella elastica* | 713.50 | Olenina, I. et al. Biovolumes and size-classes of phytoplankton in the Baltic Sea HELCOM Balt.Sea Environ. Proc. 106 (2006). |
| 118 | MMETSP0947 | Ochrophyta | *Chattonella subsalsa* | 653.00 | Domingos, P. & Menezes, M. Taxonomic remarks on planktonic phytoflagellates in a hypertrophic tropical lagoon (Brazil). *Hydrobiologia* **370**, 297-313 (1998). |
| 119 | MMETSP1160 | Ochrophyta | *Pinguiococcus pyrenoidosus* | 65.45 | Andersen, R. A., Pottert, D. & Bailey, J. C. Pinguiococcus pyrenoidosus gen. et sp. nov.(Pinguiophyceae), a new marine coccoid alga. *Phycol Res* **50.1**, 57-65 (2002). |
| 120 | MMETSP1323 | Ochrophyta | *Florenciella parvula* | 33.51 | http://roscoff-culture-collection.org/rcc-strain-details/446 |
| 121 | MMETSP1163 | Ochrophyta | *Phaeomonas parva* | 29.18 | Daiske, H. & Inouye, I. Ultrastructure and taxonomy of a marine photosynthetic stramenopile Phaeomonas parva gen. et sp. nov.(Pinguiophyceae) with emphasis on the flagellar apparatus architecture. *Phycol Res* **50.1**, 75-89 (2002). |
| 122 | MMETSP0890 | Ochrophyta | *Aureoumbra lagunensis* | 27.61 | DeYoe, H. R. *et al.* Description and characterization of the algal species Aureoumbra lagunensis gen. et sp. nov. and referral of Aureoumbra and Aureococcus to the Pelagophyceae. *J Phycol* **33**, 1042-1048, doi:DOI 10.1111/j.0022-3646.1997.01042.x (1997). |
| 123 | MMETSP0882 | Ochrophyta | *Pelagococcus subviridis* | 10.89 | Vesk, M. & Jeffrey, S. W. Ultrastructure and Pigments of 2 Strains of the Picoplanktonic Alga Pelagococcus-Subviridis (Chrysophyceae). *J Phycol* **23**, 322-336, doi:DOI 10.1111/j.1529-8817.1987.tb04141.x (1987). |
| 124 | MMETSP0167 | Rhodophyta | *Rhodella maculata* | 904.78 | http://roscoff-culture-collection.org/rcc-strain-details/655 |

This file has been converted from its original format for security purposes. Please use C56CD658F48A1 as a reference.

| | | | | | |
|---|---|---|---|---|---|
| 125 | MMETSP0011_2 | Rhodophyta | *Rhodosorus marinus* | 87.11 | Ott, F. D. Rhodosorus Marinus Geitler - a New Addition to Marine Algal Flora of Western Hemisphere. *J Phycol* **3**, 158-&, doi:DOI 10.1111/j.1529-8817.1967.tb04651.x (1967). |

This file has been converted from its original format for security purposes. Please use C56CD658F48A1 as a reference.
